# Predicting population genetic change in an experimental stochastic environment

**DOI:** 10.1101/2021.04.06.438609

**Authors:** Marie Rescan, Daphné Grulois, Enrique Ortega Aboud, Pierre de Villemereuil, Luis-Miguel Chevin

**Affiliations:** CEFE, CNRS, Université de Montpellier, Université Paul Valéry Montpellier 3, EPHE, IRD, Montpellier, France; Institut de Systématique, Evolution, Biodiversité (ISYEB), Ecole Pratique des Hautes Etudes PSL, MNHN, CNRS, Sorbonne Université, Université des Antilles, Paris, France

**Keywords:** Fluctuating selection, environmental stochasticity, population genetics, environmental tolerance curve, phenotypic plasticity

## Abstract

Most natural environments exhibit a substantial component of random variation. Such environmental noise is expected to cause random fluctuations in natural selection, affecting the predictability of evolution. But despite a long-standing theoretical interest for understanding the population genetic consequences of stochastic environments, there has been a dearth of empirical validation and estimation of the underlying parameters of this theory. Indeed, tracking the genetics of a large number of replicate lines under a controlled level of environmental stochasticity is particularly challenging. Here, we tackled this problem by resorting to an automated experimental evolution approach. We used a liquid-handling robot to expose over a hundred lines of the micro-alga *Dunaliella salina* to randomly fluctuating salinity over a continuous range, with controlled mean, variance, and autocorrelation. We then tracked the frequency of one of two competing strains through amplicon sequencing of a nuclear and choloroplastic barcode sequences. We show that the magnitude of environmental fluctuations (variance), but also their predictability (autocorrelation), have large impacts on the average selection coefficient. Furthermore, the stochastic variance in population genetic change is substantially higher in a fluctuating environment. Reaction norms of selection coefficients and growth rates of single strains against the environment captured the mean response accurately, but failed to explain the high variance induced by environmental stochasticity. Overall, our results provide exceptional insights on the prospects for understanding and predicting genetic evolution in randomly fluctuating environments.

## Introduction

To what extent is evolution predictable? This question has received considerable interest from evolutionary biologists, and has become increasingly quantitative as relevant data has accumulated. Original arguments about contingency versus necessity (Monod, 1970) or replaying the life’s tape (Gould, 1989) have been replaced by more detailed empirical and theoretical studies focusing on the rate of parallel genetic evolution in replicate lines or populations exposed to similar selective pressures (Bailey et al., 2017; Chevin, Martin, et al., 2010; Stuart et al., 2017; Tenaillon et al., 2012; Yeaman et al., 2018). Only recently has predictability in the *dynamics* (rather than outcome) of evolutionary change gained more prominence (Doebeli & Ispolatov, 2014; Nosil et al., 2018, 2020; Rego-Costa et al., 2018). This question is of crucial importance for many applications of evolution where the rate of change matters more than the end result, including evolutionary rescue, pest control, antibiotic resistance, and all contexts with strong eco-evolutionary dynamics involving a race between adaptation and population growth (Anciaux et al., 2018; Gomulkiewicz & Holt, 1995; Kingsolver & Gomulkiewicz, 2003; Marrec & Bitbol, 2020; Orr & Unckless, 2014).

Investigating predictability in evolutionary dynamics requires partitioning different possible sources of unpredictability, from measurement error to genetic drift and environmental variation (Nosil et al., 2020), which is difficult to achieve in a natural context. Experimental evolution in the laboratory is an attractive alternative, since it allows controlling some aspects of the environment to quantify their influence on evolutionary dynamics over multiple generations (Buckling et al., 2009; Elena & Lenski, 2003; Kawecki et al., 2012), and using independent replicates to assess precision limits in the measurement of key evolutionary parameters such as selection coefficients (Gallet et al., 2012). Measurements of selection in such experiments can be achieved by tracking the relative frequencies of competing genotypes over time by Illumina sequencing of a DNA barcode sequence, either originating from standing variation or engineered to this aim (BarSeq, Ba et al., 2019; Levy et al., 2015).

A major factor that may alter the predictability of evolution is temporal variation in the environment (Bell, 2010; Lenormand et al., 2009). Most natural environments exhibit random environmental fluctuations - also known as stochastic noise - characterized by their magnitude (variance) and predictability (autocorrelation, or color in power spectrum, Sabo & Post, 2008; Vasseur & Yodzis, 2004). These environmental fluctuations cause fluctuating selection at the genetic and phenotypic levels, which may reduce the predictability of evolution(Bell, 2010; de Villemereuil et al., 2020; Grant & Grant, 2002). First, unaccounted sources of environmental variability (micro-environmental variation) can increase noise in frequency dynamics, thus reducing the precision of selection estimates (Gallet et al., 2012). And second, even assuming that the environment is perfectly known at a given time, its future is uncertain if it fluctuates randomly. Environmental stochasticity thus contributes to chance in evolutionary trajectories, causing allele frequencies to undergo random walks, similarly to genetic drift caused by the finiteness of populations (Chevin, 2019; Gillespie, 1977, 1979; Kimura, 1954; Nei & Yokoyama, 1976; Otha, 1972; Wright, 1948). However stochastic evolutionary dynamics, despite being random, can still be predicted in a probabilistic sense, provided we are able to accurately model them with few parameters. This explains the popularity of diffusion approximations, where most influence of genetic drift is summarized into a single parameter, effective population size (Crow & Kimura, 1970). Whether or not the same can be done for environmental stochasticity in selection is still largely an open empirical question.

Population genetic theory of evolution in stochastic environments is usually parameterized through the distribution (mean, variance, autocorrelation) of selection coefficients over time. These are indeed the fundamental parameters that drive the evolutionary dynamics and determine long term outcomes, such as probabilities of fixation (Kimura, 1954; Otha, 1972) or expected heterozygosities (Nei & Yokoyama, 1976). However in most empirical contexts, it should also be of interest to project evolutionary change based on how *the environment itself* fluctuates. Global and local time series of environmental variables are available from sources such as the IPCC, and are also straightforward to obtain anew using automated devices such as thermochrons. The challenge is then to go from variation in the environment to variation in selection, to respond to quantitative questions such as: How much variance in selection is caused by variance in the environment? Does environmental variation affect the mean selection coefficient? And is there an influence of environmental autocorrelation − which determines the predictability of environmental fluctuations − on patterns of fluctuating selection?

Establishing such a link between fluctuations in the environment and in selection is not straightforward. Selection arises from the differential growths of genotypes in competition, and its variation in fluctuating environments can have different sources, which need to be deciphered in order to improve predictive power. For instance, if selection at any time point only depends on the current environment a population is experiencing, then selection coefficients may be measured at only a few constant values of the environment to produce a type of “selection reaction norm”, which may then be combined with the pattern of environmental fluctuations to project population genetic change. If applicable, this would drastically reduce the sequencing effort involved in estimating selection and predicting evolutionary change in a stochastic environment. The prospects for prediction would further be spectacularly improved if the growth rates of genotypes in competition (and hence selection) could be predicted from their growth rates in monculture across environments. The latter is described as an environmental tolerance curve (Lande, 2014; Lynch & Gabriel, 1987; Rescan et al., 2020), and is straightforward to measure, requiring no sequencing effort (since genotypes are grown in monoculture). Such simple mechanistic links between the environment and selection, beyond just facilitating projections, may also allow reaching a better understanding of environmental variation in selection, which may prove useful when extrapolating to other environmental regimes, or even organisms. However, the usefulness of such mechanistic approach first needs to be evaluated in controlled conditions, to assess the limits of its underlying approach. For instance, the selection reaction norm approach would be compromised if selection at a given time depends on the sequence of environments a population was exposed to, because of a memory of past environments mediated by phenotypic plasticity (Botero et al., 2015; Rescan et al., 2020). Similarly, fitness in competition may depend on specific interactions between genotypes, causing selection to be frequency- or density-dependent (Chevin, 2011), and precluding its accurate prediction from growth in monoculture.

We here used experimental evolution in the laboratory to investigate population genetic changes caused by a randomly fluctuating environment. We worked with the halotolerant micro-alga *Dunaliella salina*, which can thrive across a broad range of salinities. Contrary to previous experiments that also focused on random variation in the environment (Boyer et al., 2021; Dey et al., 2016; Wieczynski et al., 2018), we here allowed salinity to vary over a continuous range (rather than switching randomly from low to high), with a controlled distribution over time. We used a liquid-handling robot to control the mean, variance, and autocorrelation of salinity, and tracked the frequency of standing genetic variants through time by Illumina amplicon sequencing. We have previously shown that the stochastic demography of these populations was well predicted by combining patterns of environmental variation with short-term salinity tolerance curves (Rescan et al., 2020). Here, we ask how key parameters that appear in classic models of stochastic fluctuating selection are influenced by parameters of environmental variation, and whether population genetics in a stochastic environment can be predicted from first principles.

## Results

### Tracking population genetics in an experimental stochastic environment

We followed the frequency of one strain (CCAP 19/15, hereafter denoted as C) of the microalgae *Dunaliella salina* competing in a mixture with another strain (CCAP 19/12, hereafter denoted as A) during 37 transfers (∼100 generations), in constant or randomly varying salinity. We transferred each line twice a week (every 3 or 4 days), diluting 15% of the population of origin into fresh medium with controlled salinity, which differed across lines. Our treatments consisted of 138 independent fluctuating salinity time series with the same mean (*μ_E_* = 2.4M NaCl) and variance 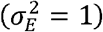 but four autocorrelation levels, from negative (*ρ* = –0.5) to highly positive (*ρ* = 0.9) (insets in Figure 1a-d). We also had three constant treatments with salinity fixed to 0.8, 2.4 or 3.2M NaCl (insets in Figure 1e), with 4 to 5 replicates for each constant salinity. We used Illumina amplicon sequencing of a nuclear and a choloroplastic marker to track the frequency dynamics of strain C within and across lines under these different treatments (Figure 1). Some lines went extinct over the course of the experiment (Rescan et al., 2020), so the sample size over which we investigated fluctuating selection decreased over time (numbers next to the curves in **Figure 1**).

We analyzed how the mean, variance and autocorrelation of salinity influence strain frequency dynamics using a state-space logistic regression model. Briefly, this model assumes that the observed strain frequency is binomially distributed, while the true logit allelic frequency *ψ* follows a Gaussian process over time, with mean 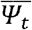 variance (*ψ_t_*) at time *t* provided by

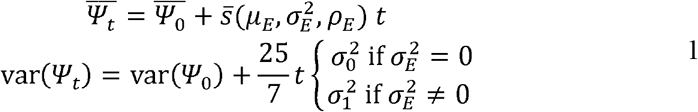

**Figure 1:**
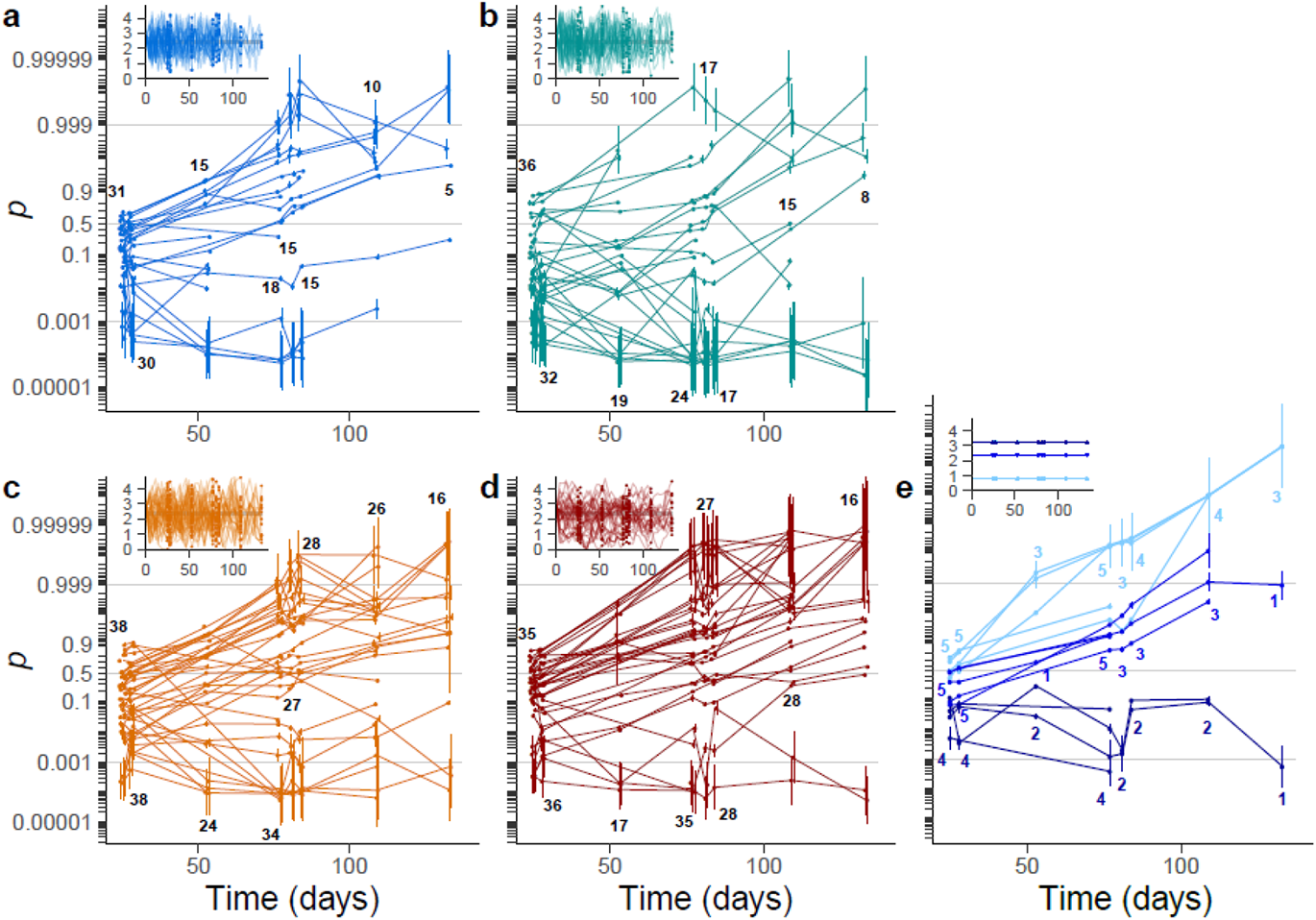
Frequency dynamics in fluctuating versus constant environments. Logit frequencies of all lines in fluctuating (from blue to red: autocorrelation −0.5 (a), 0 (b), 0.5 (c), 0.9 (d)) and constant (e) salinities (from lighter to darker: 0.8M, 2.4M and 3.2M) are represented, with the corresponding salinity time series plotted in the insets. Frequencies were estimated as random parameters in the state-space logistic regression of choloroplastic and ITS2 marker sequences. Each line represents the frequency dynamics in one population, and numbers are the total number of populations sequenced in each salinity treatment and at each time point.

The mean selection coefficient 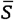 is thus assumed to depend on the mean *μ_E_*, variance 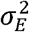 and autocorrelation *ρ_E_* of the environment. The variance in selection coefficients is 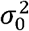 or 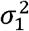 in constant vs fluctuating salinity (environmental variance 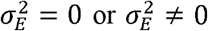), respectively, while the coefficients 25/7 corrects for the fact that salinity constant for a few days in between each transfer (Appendix; more detail on the model is provided in the Methods).

#### The average selection is reduced in fluctuating environments

The parameters of environmental fluctuations had a strong impact on the dynamics of population genetic change (as summarized in Table 1). The mean dynamics of logit-frequency (and therefore the mean selection coefficient 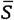) across replicates depended on the mean salinity, with larger positive selection for strain C at lower salinity (P < 10^−34^, Table 1). In addition, the mean selection coefficient was also strongly influenced by the temporal variance in salinity (P < 10^−20^), with lower advantage for strain C in stochastic environments, as compared to constant environments with the same mean (2.4M NaCl; compare black and middle blue line in **Figure 2.a**).

**Figure 2:**
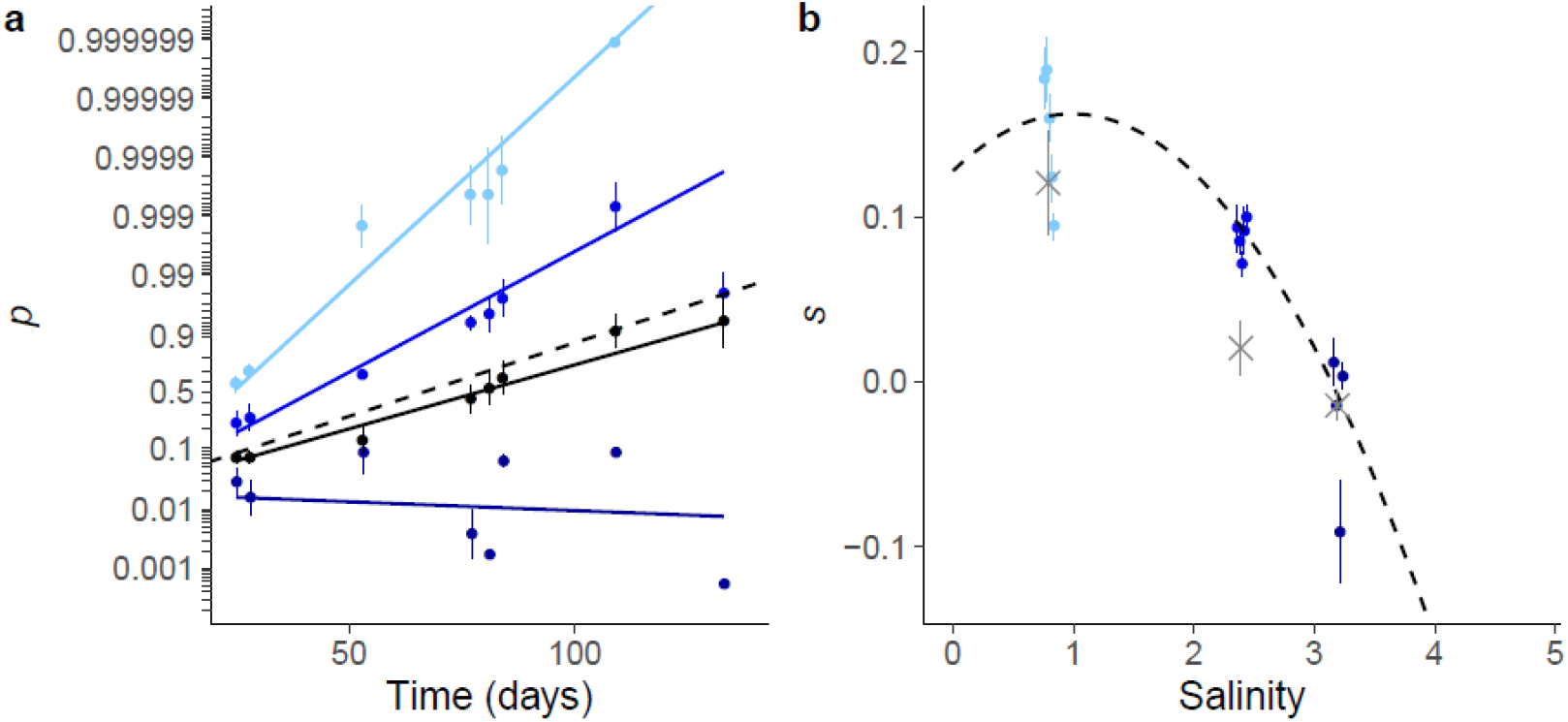
Mean selection in fluctuating versus constant environment. (a) Mean trajectories of logit allelic frequency over time. Solid lines represent logistic regression fits (eqs. (1), and (2)), while dots and error bars are time-specific estimates and bootstrapped standard deviations of logit frequencies, based on random factors in the state-space model. Black: fluctuating salinity. Blue: constant salinities, from lighter to darker: 0.8M, 2.4M and 3.2M. The dashed line is the mean logit frequency dynamics as predicted from the selection reaction norm in panel b. (b) Selection reaction norm in constant environment. The selection coefficient for strain C is shown as dots and error bars for the logistic regression fit independently in each population (colors match those in panel a). The dashed line represents the model that includes the influence of salinity on selection (selection reaction norm), and the gray crosses are the prediction from the difference in growth rates between strains A and C (with error bars corresponding to the standard error).

**Table 1:** Effect of the temporal mean *μ_E_*, variance 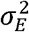 and autocorrelation *ρ_E_* of the environment on the mean (Fixed effets) and variance (Random effects) of strain C frequency (Estimates, Standard errors and P-values from Wald test). All terms representing an interaction with time *t* measure an effect of the environment on selection (letters correspond to coefficients in eq. (11), while 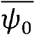 and *V*(*ψ*_0_) are the mean and variance of initial strain C frequency.

| Fixed effects : $\overline{\psi}_t$ | | | |
| --- | --- | --- | --- |
|  | Estimate | Std. Error | Pr(>\|W) |
| $t (\theta_0)$ | 0.0925 | 0.00794 | 2.34E-31 |
| $t: \mu_E (\theta_1)$ | -0.0982 | 0.00795 | 4.77E-35 |
| $t: \mu_E^2 (\theta_2)$ | -0.0349 | 0.00689 | 4.10E-07 |
| $t: \sigma_E^2 (\theta_3)$ | -0.0776 | 0.00845 | 3.98E-20 |
| $t: \rho_E^2 (\theta_4)$ | 0.0900 | 0.0109 | 1.82E-16 |
| $\overline{\psi}_0$ | -3.82 | 0.266 | 1.12E-46 |
| Random Effect: $V(\psi_t)$ | | | |
| $V(\psi_0)$ | 0.0344 | 0.107 | 7.46E-01 |
| $t: [\sigma_E^2 = 0](\sigma_0^2)$ | 0.0186 | 0.00405 | 4.32E-06 |
| $t: [\sigma_E^2 = 1](\sigma_1^2)$ | 0.131 | 0.00845 | 1.76E-54 |

To investigate to what extent this effect of environmental stochasticity on the mean coefficient selection can be predicted from first principles, we measured the direct influence of the environment on selection using a selection reaction norm in constant salinity (**Figure 2.b**). We assumed a quadratic shape for selection as function of a constant environment,

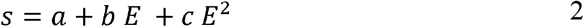

where *E* is the (constant) environment, measured as deviation from the mean salinity. We found that in a constant environment, strain C is favored in the intermediate environment where salinity is 2.4M NaCl (*a* > 0, p < 10^−31^, see Table 1), but is less favored at higher salinity (*b* < 0, P < 10^−34^), with an advantage that vanishes towards 3M NaCl. In addition, there is a negative quadratic effect of constant salinity on selection (*c* < 0, P < 10^−6^), such that the salinity reaction norm is concave, with an optimum at an intermediate salinity (**Figure 2.b**). For the reaction norm estimated by eq. (2), the maximal selection coefficient for strain C is 0.16, which occurs at an optimal salinity of 1.0M, and the breadth of the hump around the optimum (defined as 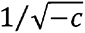) is 5.5M (**Figure 2.b**).

Assuming that selection is density- and frequency-independent, and is not affected by any memory of past environments (Rescan et al 2020, and below), then selection coefficients measured over a range of constant environments (selection reaction norm) can be used to predict the dynamics of (logit) allele frequency in a fluctuating environment, by summing selection over all past environments experienced by both strains (Chevin 2011, 2019). Combining the reaction norm in eq. (2) with the (nearly) normal distribution of salinities in our evolutionary experiment, the predicted mean selection coefficient is

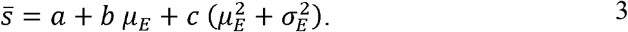

Eq. (3) highlights that in a stationary stochastic environment, environmental variance affects the mean selection coefficient only if there is a quadratic effect of salinity on selection (*c* ≠ 0). In contrast, when *c* = 0 eq. (3) predicts the same expected (logit) frequency dynamics in constant vs stochastic environments, if they have the same mean *μ_E_*. The expected logit strain mean frequency 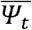 over time based on the quadratic selection reaction norm (eq. 3) is shown as dashed lines in **Figure 2.a**, and matches very well the mean frequency dynamics estimated in fluctuating environments (solid black line in **Figure 2.a**; P = 0.28 for the Wald test between selection estimate and reaction norm prediction). This indicates that the reduced expected selection coefficient in a fluctuating environment likely arises from the concavity of the relationship between selection and the environment.

### Environmental predictability also altered the expected frequency dynamics

Patterns of stochastic environmental fluctuations are not only characterized by their variance, which relates to their magnitude, but also by their autocorrelation, which determines how salinity transitions take place. This can have a major impact on the population dynamics of each strain in isolation in this species (Rescan et al., 2020), and may therefore also alter selection in a stochastic environment. We found that autocorrelation had a substantial influence on mean selection (P < 10^−15^, LR test between the regression described by eq. (12) and the same model without autocorrelation effect on mean selection). The mean selection for strain C in stochastic environments significantly increased with increasing squared autocorrelation 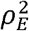 of salinity (Figure 3). The predictions from eq. (3) matched well the dynamics under intermediate predictability (autocorrelation *ρ* = ±0.5, orange and blue dots and lines, compared to prediction in dashed black line in **Figure 3**), while in the more autocorrelated environment where transitions are smallest (*ρ* = 0.9), the mean selection coefficient exceeded the predictions from eq. (3), and was instead very close to that under constant salinity 2.4M (the mean of the fluctuating treatments; compare red and gray dots and lines in **Figure 3**). Importantly, we found no significant difference between autocorrelation 0.5 and −0.5 (P = 0.45, likelihood ratio test between and the model described by eq. (1) and the full model that also includes an effect of unsquared autocorrelation *ρ_E_*). This indicates that allele frequency dynamics did not respond to the magnitude of salinity transitions, controlled by their temporal autocorrelation, but rather to their predictability, as determined by the squared autocorrelation.

Because temporal autocorrelation does not change the stationary distribution of environmental values, but only the similarity between two successive environments, the significant effect of environmental predictability must involve an effect of past environment on current selection. In fact the salinity reaction norm in eq. (2), which only depends on current environment, predicts no influence of environmental autocorrelation on mean selection (dashed line in Figure 3b). In order to capture the influence of past environment on selection, we used the known past and current salinities *E*_t-1_ and *E*_t_ from successive transfers (at steps 6-7, 20-21 and 21-22 of the experiment) as predictors for selection in the experiment with fluctuating salinity. Within the state-space framework with observations described by eq. (1), this led to

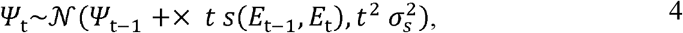

where *Ψt*–1 and *Ψ_t_*are the logit frequency before and after each transfer (over a short duration of *t* = 3 or 4 days), and *s*(*E*_t-1_, *E*_t_) models a bivariate selection reaction norm in response to the current and previous salinity (see Supplementary Table 1):

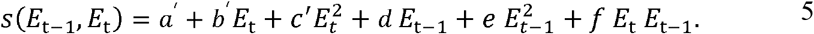

**Figure 3:**
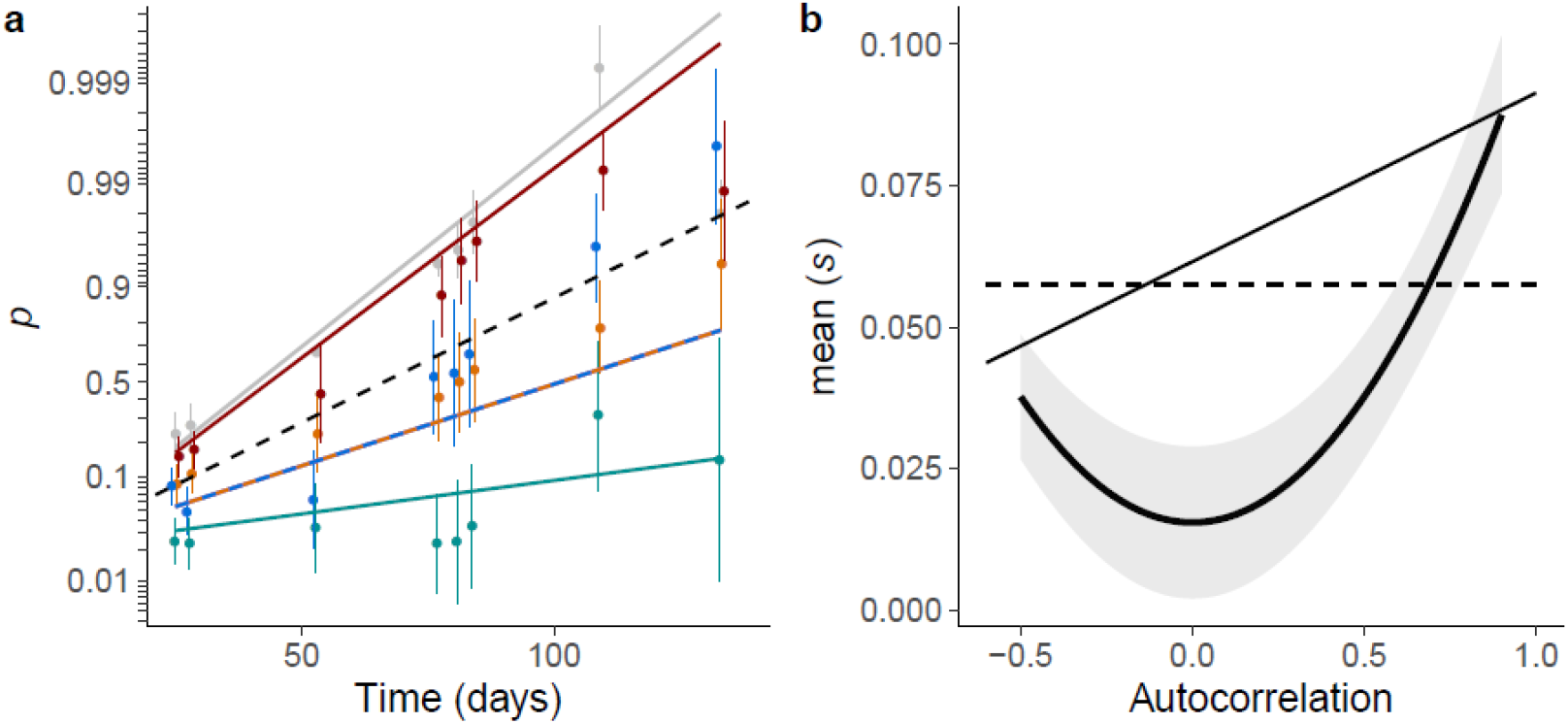
Influence of environmental autocorrelation on selection. (a) Frequency dynamics for treatments with different autocorrelations (From blue to red: −0.5, 0, 0.5 and 0.9). Dots and error bars are the mean and standard errors bootstrapped from the realized logit frequencies, and lines give the logistic state-space model predictions (eqs. (1) and (2)). Gray: constant environment with same salinity mean 2.4M. Dashed black line: prediction from the selection reaction norm built in constant environments (eq. ()). (b) Selection response to autocorrelation. The black thick curve and ribbon show the mean selection coefficient estimated by the logistic state-space regression and the 95% confidence intervals computed with the delta method. Lines are the predictions from the univariate (dashed line, eq. ) and bivariate (solid line, eq. ) selection reaction norm.

We found a significant effect of current (P = 2.1 10^−2^, LR test between a model considering only current salinities and a model with constant selection) and past salinities (P = 5.6 10^−3^, LR test between a model with a bivariate vs a univariate selection reaction norm) on selection coefficients. (We additionally injected terms for the influences of population density and strain frequency-dependent in eq. (5), but found no signal of density- (P = 0.51) nor frequency-dependent selection (P = 0.09).) Combining with the binormal distribution of salinities at two successive transfers, we found that the predicted mean selection coefficient was higher in more autocorrelated environment as observed in our experiment, but was largely overestimated in unpredictable environments with *ρ = 0* (thin line vs thick line and ribbon in **Figure 3b**).

### Selection reaction norm under-predicts selection variance

The variance of allele C frequency substantially increased in a stochastic environment (P < 10^−33^, **Figure 4.b**), consistent with predictions from theoretical population genetics (Gillespie, 1973, 1991; Kimura, 1954; Otha, 1972; Wright, 1948). The variance in selection 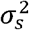 was over seven times higher in stochastic as compared to ‘constant treatments. We nevertheless observed a significant increase in allele frequency variance 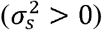 even in constant salinities, which may result from genetic drift and/or from micro-environmental variation impossible to control for (slight light and temperature heterogeneity in the incubator).

**Figure 4:**
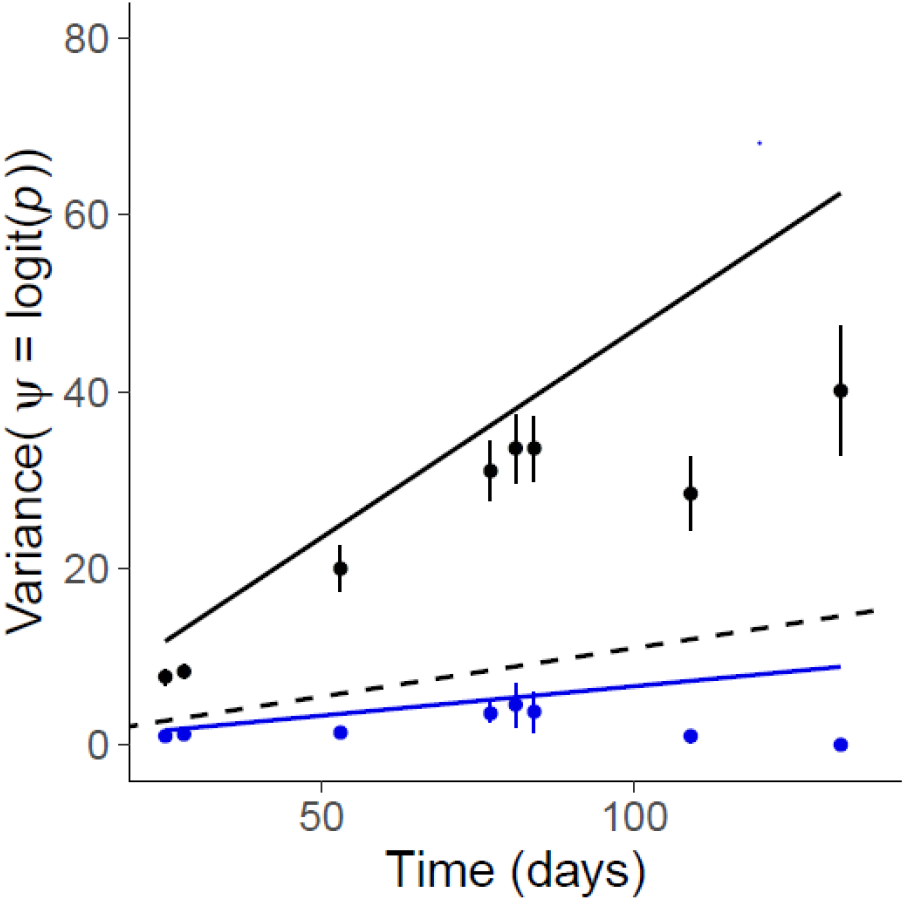
Frequency variance in constant versus fluctuating environments. Solid lines represent the logistic regression fit (eqs. (1)) in constant (blue) and stochastic (black) environments, while the dashed line is the prediction from the univariate selection reaction norm (eq. (2)). Dots and error bars are the mean and bootstrapped standard deviation of the realized logit frequencies estimated as random factors in the state-space model. Variance was the same in all constant treatments (blue) and all fluctuating treatments (black), and bootstraps were performed after subtracting the mean frequency.

From the univariate selection reaction norm in eq. (2), the expected variance of selection coefficients caused by fluctuating selection is:

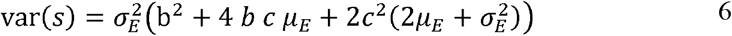

This predicts a selection variance 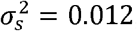, which is about one order of magnitude below the variance measured in fluctuating environment (0.13). Selection variance given by the logistic regression was the sum of selection variance linked to salinity fluctuations, predicted by eq. (3), plus a residual variance corresponding to the measure in a constant environments (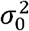 in eq. (1)). We injected this predicted variance in eq. (1) to compute the expected variance of logit transformed allelic frequency. However, even after adding the selection variance measured in constant environments, predictions from the selection reaction norm remained more than 4 times lower than the variance estimated in the logistic regression (**Figure 4**).

Eq. (6) predicts that the variance in selection coefficients increases with the environmental variance, but is not affected by autocorrelation, if selection only depends on the current environment. We obtained the selection variance predicted by the bivariate selection reaction norm in eq. (5), which also includes an influence of the previous environment, but this barely changed the prediction for selection variance, which was still drastically below the variance estimated in constant environments (between 0.025 and 0.03 when summed to the selection variance in constant environment).

### Tolerance curves accurately predict selection only in constant environments

In a population comprising two genotypes, the selection coefficient can be translated into a difference between the *per-capita* population growth rates of genotypes in competition (Chevin, 2011),

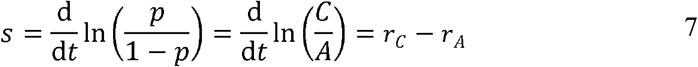

where and *C* and *A* are the population densities of the two genotypes, and *r_C_* and *r_A_* their *per-capita* growth rates (equal to the rate of change on the log scale). If selection is density-independent (meaning that the *per-capita* growth rates of both genotypes respond identically to population density), and there are no interactions between strains (no frequency dependence), then the intrinsic growth rates of both strains, each measured in monoculture, can directly be used to predict the outcome of selection in a given environment (Chevin, 2011). In addition, selection in a fluctuating environment can be then predicted by comparing the tolerance curves of both strains, which describes how their intrinsic rate of increase measured in monoculture changes with the environment (**Figure 5**). In particular, genotypes with the same tolerance breadth would then be expected to have a linear selection reaction norm, leading to a normal distribution of selection coefficients if the environment follows a Gaussian process (Chevin, 2019). Conversely, any curvature in the selection reaction norm in eq. (2) would be interpreted as a difference in tolerance breadths of the strains, reflecting differences in the plasticity of underlying traits influencing fitness across environments (Chevin, 2019; Chevin & Lande, 2010), and leading to skewed distribution of selection coefficients (**Figure 5**).

**Figure 5:**
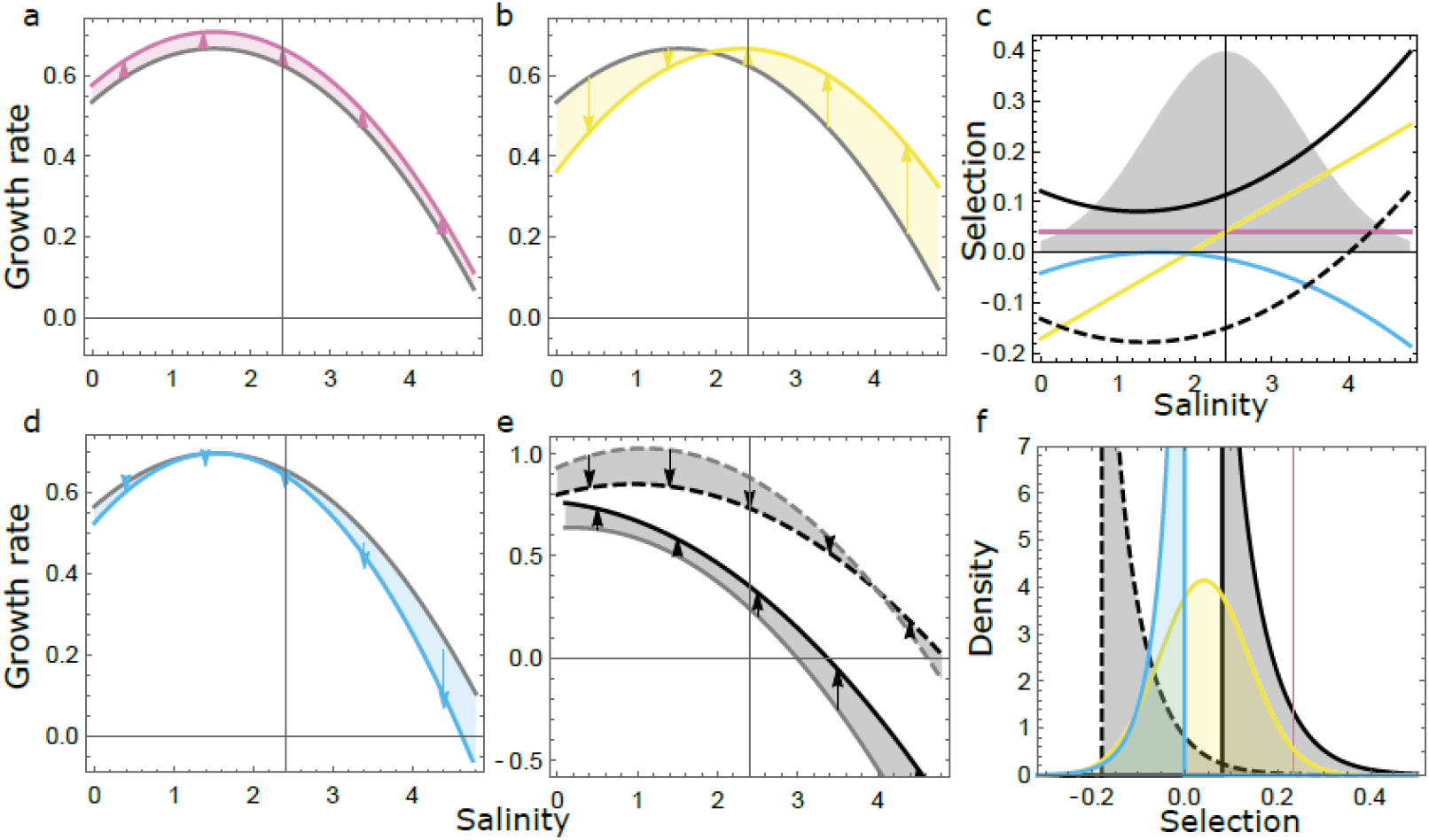
Relationship between tolerance curves and selection across environments. Panels a-e represent tolerance curves, with absolute fitness (population growth) in isolation plotted against the environment (here, salinity). In panels a, b and d, the tolerance curve of a reference strain (in gray) is contrasted to that of an alternative genotype or mutant (in color), which varies in only one parameter of the tolerance curve: maximal growth (a), salinity optimum (b), or niche width/breadth (d). Selection for the mutant equals the difference in growth rates between strains (shown as arrows) if fitness is density- and frequency-independent. In addition, the past environment may influence the current tolerance curve of each strain, as illustrated in panel e, where the tolerance curve of the mutant (in black) and reference strain (in gray) vary depending on whether they were transferred from low (plain line, 0.5M) or high (dashed, 4M) salinity. Panel c shows the selection reaction norm for the mutant based on these tolerance curves (hence assuming density- and frequency-independent selection), using the same line types and colors as in panels a,b, d & e. The gray normal curve materializes the salinity distribution used in our fluctuating salinity experiment (rescaled vertically for graphical convenience). Panel f plots the resulting distribution of selection coefficients in the fluctuating environment, with line types and colors as in previous panels. Note that the distribution of selection coefficient is Gaussian if both strains have the same tolerance breadth (yellow), but can otherwise be highly skewed.

In our fluctuating treatments, there was no correlation between selection measured over one salinity transfer and strain differences in per-capita population growth rates, as estimated by Rescan et al. (2020) (Supplementary Figure 2). This is not necessarily surprising, given that most variance in our stochastic treatments was not explained by our salinity tolerance curves. By contrast, selection estimated over 37 transfers in constant environments was in a good agreement with differences between the per-capita growth rates of strains in monoculture (non-significantly different at salinity = 0.8 and 3.2M, but slightly below the selection measured in competition at 2.4M, P <10^−6^, Wald test; see gray crosses in **Figure 4**). A notable discrepancy was that the growth rates in monoculture predicted a linear selection reaction norm, because the two strains have similar salinity tolerance breadths (95% bootstrapped confidence interval: [-6 10^−3^, 2 10^−2^], from 1000 simulations and fit of a quadratic reaction norm). Based on this, we would thus predict that the mean allelic frequency does not differ between stochastic and constant environments (eq. 3), contrary to what we observed in our experiment.

## Discussion

The extent to which evolution in a stochastic environment can be predicted is still largely an open question. The intense selectionist vs neutralist debate in the second half of the XXth century somewhat crystallized around the relative roles of drift vs fluctuating selection as sources of chance in population genetics (Fisher & Ford, 1947; Gillespie, 1977; M, 1968; Yamakasi & Maruyama, 1972). This question finds ramifications in nowadays’s science (Miura et al., 2013), but its answer ultimately rests on empirical quantitative evidence: it depends on how the variance in selection compares to the reciprocal of the effective population size. Using over 150 replicate lines of two competing strains exposed to fluctuating salinities with controlled mean, variance, and autocorrelation, we experimentally quantified the influence of random environmental fluctuations on stochasticity in selection. We found that selection variance was seven times higher in stochastic than in constant salinities, and that environmental fluctuations also impacted the mean selection coefficient, in ways that could partly be predicted from selection and growth in constant environments.

One of the emerging debates in the early literature on fluctuating selection regarded whether or not stochastic fluctuations influence the expected strength of selection, and mean evolutionary trajectory. Some models found that variance in selection coefficients influences the mean trajectory (Gillespie, 1973; Wright, 1948), and others that it does not (Kimura, 1962; Nei & Yokoyama, 1976), depending notably on the role of density regulation in selection (Nei, 1971; Nei & Yokoyama, 1976). More recent work has re-explored this question from another standpoint, by explicitly including a phenotype under selection for a moving optimum, and predicted that the variance of fluctuations should only influence the expected selection if genotypes differ in their phenotypic plasticity and environmental tolerance (Chevin, 2019). Here, we found that the mean selection coefficient favoring strain C is substantially lower in a randomly fluctuating environment as compared to a constant environment with salinity fixed to the mean of the fluctuating treatment, as expected if the two strains differ in their plasticity levels (Chevin, 2019). This is also consistent with the concave shape of the selection reaction norm measured in constant environments, suggesting that our focal strain C has a narrower tolerance niche (Chevin, Lande, et al., 2010).

In addition to the amplitude of environmental fluctuations, their autocorrelation pattern also affected the mean frequency trajectory, with higher selection for our focal strain in the more predictable environments (Figure 3). This suggests that selection depends not only on the current environment, but also on environmental transitions. Accordingly, taking the previous salinity into account improved the prediction for the expected frequency dynamics in temporally autocorrelated stochastic environment. This is consistent with the crucial effect of past salinity on population growth in *Dunaliella salina*, involving a transgenerational phenotypic memory mediated by the dynamics of glycerol content as osmoprotectant (Rescan et al., 2020).

The frequency dynamics in fluctuating vs constant environment, and the selection reaction norm measured in constant environments, both point towards strain C being less plastic than its competitor. Finding that it is more advantaged in more predictable environments (Figure 3) therefore seems at odds with the theoretical prediction that higher levels of plasticity should be favored in more predictable environments (Botero et al., 2015; De Jong, 1999; Gavrilets & Scheiner, 1993; Lande, 2009, 2014; Tufto, 2015). Furthermore, we found in a previous study that followed up on this experiment that higher levels of morphological plasticity were maintained in pure lines that evolved in highly predictable environments (Leung et al., 2020), consistent with theoretical predictions. Despite this discrepancy, both studies share striking features. They both revealed (i) an evolutionary outcome that is very similar between constant and highly predictable fluctuating environments, and (ii) a response that depends on environmental predictability (squared autocorrelation) rather than on temporal autocorrelation *per se*, with similar dynamics in negative and positive autocorrelation with the same absolute value. The latter is not trivially explained by the effect of salinity transitions on fitness, because the selection reaction norm with effects of past and current environments predicts a linear response of mean selection to autocorrelation (**Figure 3.b**; see also Rescan 2020 for the effect on growth rates of individual strains). As temperature in most continents have become bluer – more negatively autocorrelated − over the past century (García-Carreras & Reuman, 2011), there is a growing need to understand how environmental time series with opposite autocorrelation can lead to similar genetic dynamics. Possible explanations may involve a more complex shape of the selection reaction norm (e.g. involving higher order terms such as 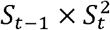, which measures the effect of past salinity on the breadth of the selection reaction norm), or an effect of the salinity two transfers in the past, with correlation *ρ*^2^ to the current salinity. Testing such hypotheses would require challengingly long time series of allelic frequency. On the other hand, another line of evidence may come from an analysis of the molecular mechanisms involved, such as transcriptomic and epigenomic responses to salinity transitions in lines that evolved under different autocorrelation treatments (Leung et al 2020, *in prep*).

Our study underlines the difficulty in predicting population genetic change in stochastic environments from more basic (and less labor intensive) data, such as tolerance curves or selection reaction norms measured in constant environments. Data on growth rates of individual genotypes are the easiest to obtain for clonally reproducing individuals, because they only require counts and do not rely on sequencing effort. However here, they only gave a qualitatively acceptable prediction for selection, and only when measured in constant environment and in the long run. This points to interactions between strains causing their growth rate in competition to differ from that in monoculture, yet we did not detect any evidence of frequency-dependent selection. Another challenge is that a substantial part of the variance in allelic frequency change was not explained by simple responses to salinity captured by our selection reaction norms (Figure 4). In addition, we found significant variance in frequency change also in constant environments, pointing to uncontrolled sources of stochasticity even in laboratory conditions, possibly involving micro-environmental variation or drift due to the random sampling of individuals (for instance because of bottlenecks upon transfers, Wahl et al., 2002). Overall, our results thus indicate that randomly fluctuating environments can strongly shape the dynamics of population genetic change, but that deciphering and predicting these effects may require more detailed information than is provided by population growth or even selection in a constant environments.

## Methods

### Evolutionary experiment

We exposed a mixture of two *Dunaliella salina* strains to fluctuating vs constant salinity, and tracked their frequencies through time by amplicon sequencing. Details of this long-term experiment were described in a previous study focused on demography (Rescan et al., 2020), so we only summarize them briefly here. Populations of *Dunaliella salina* were initiated by mixing 50% of strain CCAP19/12 (denoted below as strain A) and 50% of strain CCAP19/15 (hereafter strain C), and exposed to constant or fluctuating salinities during 37 transfers (∼100 generations). Populations were transferred twice a week by diluting 15% of the culture into 800 µL of fresh medium using a liquid-handling robot (Biomek NXP Span-8; Beckman Coulter). At each transfer, the target salinity was achieved by mixing the required volumes of hypo- ([NaCl] = 0 M) and hyper- ([NaCl] = 4.8 M) saline media, after accounting for dilution of the pre-transfer salinity. Populations in constant salinities were exposed to 0.8, 2.4 and 3.2M NaCl, with 5 replicates per salinity, while the fluctuating treatments consisted of 39 independent stochastic salinity time series (with the first replicated three times), for each of 4 temporal autocorrelation levels. Salinities were sampled from a first-order autoregressive process (AR1) with mean 2.4M, variance 1 and autocorrelation −0.5, 0, 0.5 and 0.9.

Populations were frozen after transfers 6, 7, 8, 14, 21, 22, 23, 30 and 37 for extraction and amplification of a chloroplast locus and the ITS2. Before extraction, *Dunaliella* cells were killed by adding 120 µL of 100% ethanol to the 480 µL of culture left for each populations after replication and demographic measures (Rescan et al., 2020). After careful mixing, plates were centrifuged 5 min at 6000 rpm, supernatant removed, 200 µL PBS added for conservation and samples resupended. Plates were frozen at −20°C until extraction.

### Data acquisition

#### Sequencing

Genomic DNA was extracted using the Nucleospin® plant II (Macherey-Nagel) following the manufacturer’s protocols. Population density varied along time and between lines in our experiment (Rescan et al., 2020), and only the 1015 samples with cell density greater than 10^3^/mL were extracted (representing 88% of all AC populations that were not extinct by the time of the transfer).

All samples were amplified for the ITS2 segment of the ribosomal gene, and a chloroplast locus. The primer of ITS2 gene amplified a fragment of 200bp, by ITS2-for2 (^5’^-GCAGAATTCCGTGAATCATCAAATC-^3’^) and ITS2-rev2 (^5’^-GCGAGCGATAAGCTGCCTACCCAGTTG-^3’^). For the chloroplast locus, new primers specific of each strain were drawn from Whole Genome sequencing of the strains (unpublished data), in order to obtain one fragment of 200bp. The primer of the chloroplast locus amplified a fragment of 200bp, by Chloro53-for1 (^5’^-CGTTTATCCATATACGGG-^3’^) and Chloro53-rev2 (^5’^-CGCGCGAGTACCATCAGGACC-^3’^). Both loci were amplified separately for each sample, with their specific primers and the addition of specific sequences to anchor indexing primers: forward (^5’^TCGTCGG-CAGCGTCAGATGTGTATAAGAGACAGYR- [primers of ITS 2]-^3’^) and reverse ^5’^GTCTCGTGGGCTCGGAGATGTGTATAAGAGACAGYR- [primers of ITS 2]-^3’^) for the ITS2, and Chloro-N (^5’^TCGTCGGCAGCGTC AGATGTGTATAAGAGACAGN- and - NN - and - NNN - [primer Chloro]^3’^) and reverse (^5’^GTCTCGTGGGCTCGGAGATGTGTATAAGAGACAGN - and - NN - and - NNN - [primer of Chloro]^3’^) for the chloroplast locus. PCRs were carried out in a final volume of 20 μl containing 0.05-0.1 µg/ml template DNA, Phusion High-Fidelity PCR Master Mix with HF Buffer 2X (ThermoScientific), 1 µM forward and reverse primers. PCR amplifications were conducted in Eppendorf Mastercycler ep gradients S thermal cycler under the following conditions: 1 min initial denaturation at 94°C, 35 cycles of 1 min at 94°C, 1 min at T_m_°C, 1 min at 72°C, and a final extension at 72°C for 5 min. The annealing temperatures (T_m_°C) were 55°C for the ITS2 and 70°C for the chloroplast locus. Negative controls were included in both the extraction and amplification steps. All amplicons were checked on the agarose gels (1%) and 901 positive samples (for both the ITS2 and the chloroplast locus) were send to the sequencer.

Sequencing was performed by the “Genseq” platform of LaBEX CEMEB, Montpellier. For each sample, PCR products for the ITS2 and chloroplast locus were mixed in the same tube in balanced quantity, and purified. All samples underwent an additional step of amplification with specific primers that included 9-bp long tags, to construct libraries for Illumina sequencing. Libraries were normalized, pooled, and sequenced on 1 run of the MiSeq Sequencer (Illumina, San Diego, CA) using the 2 × 150 bp MiSeq Reagent Kit v2.

The quality of reads was satisfying, with only few reads having quality below 20 (phred score, FastQC). We obtained 1718 demultiplexed paired-end fastq files, corresponding to the 859 samples. Fastq files of the chloroplast locus and ITS2 sequences were split with a bash script matching the ITS2 and chloroplast locus primers. Only exact matches where retained, rising the dataset quality. This produced unpaired reads that were repaired using repair.sh from BBMap 38.3254. Reads shorter than the length of the ITS2 segment were removed using trimmomatic55. Reads were finally merged using bbmerge-auto.sh from BBMap, with more than 90% of the reads merging successfully for all samples. The output quality was re-analyzed with FastQC and was satisfactory (phred score > 30). Samples with no reads left were removed, leaving 853 ITS2 and 842 chloroplast locus files, with an average of respectively 3407 and 1422 reads.

#### Haplotype tagging

Importantly, strains A and C are not isogenic, so they may have shared variants. In addition, most of the sequences were present in only one or few copies due to sequencing errors or punctual mutations. In order to make efficient use of the sequencing data for tracking the frequency of strain C in a mixture with strain A, we reduced all chloroplast and ITS2 reads to short haplotypes made of a succession of few linked SNPs that individually maximized the *F_ST_* among pure, reference cultures. In addition to strains A and C, we consider *Dunaliella salina* strain CCAP 19/18 used in the same experimental set up for another study (Rescan et al., 2020), to be able detect potential cross contamination. For each polymorphic site, we computed the global Nei *G_st_* for multiple alleles (Nei, 1973):

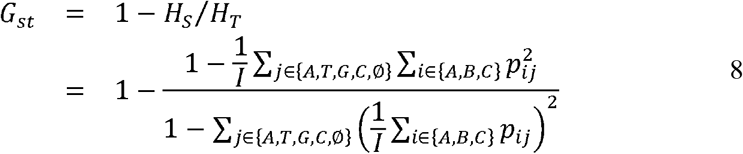

where *H_S_* is the mean expected heterozygosity within strain (that is, the probability that two haplotypes drawn from the same strain are different), *H_T_* the expected total heterozygosity (probability that two haplotypes drawn from the full sample combining all strains are different), *p_ij_* the frequency of base *j* (possibly A, T, G, C or ø) in strain *i*, and *I* = 3 the total number of strains. For the chloroplast locus, we built haplotypes by keeping all sites with *G_ST_ >* 0.8. All haplotypes from the pure strains A and C were specific to that strain, consistent with our design of strain-specific markers. More than 99.9% of the reads in the experimental population matched a reference haplotype and were tagged accordingly allele A or C. The remaining reads were removed in subsequent analyses. For the ITS2 locus, strains A, B and C were much less differentiated, and haplotypes were built using 3 bases displaying a G_ST_ > 0.2. We first noticed the presence of *Dunaliella viridis* ITS2 alleles (about 31% of the total number of ITS2 sequences in pure strains A and C), evidencing a contamination of our references strains. Such contamination was absent in all experimental populations, indicating either that contamination occurred after the end of the experiment (the reference populations were extracted 16 months afterwards), or that *D. viridis*, if initially present, disappeared rapidly before the 7^th^ transfer in our experiment. In our experimental AC mixed populations, more than 99.9% of the reads matched two ITS2 reference haplotypes, and the remaining ones were removed. The first haplotype was specific to strain A (and was thus tagged as A allele), while the other one was shared by both strains but more common in strain C (present in 100% of the reference strain C, but only about 20% of reference strain A), so it was tagged as C allele.

#### Calibration

To validate and calibrate our estimates of strain frequencies based on relative number of Illumina reads, we mixed references cultures of strains A and C in 800 µL, with predefined relative frequencies 0, 5, 10, 20, 30, 40, 50, 60, 70, 80, 90 and 95 and 100% (where 0 and 100% are the pure cultures described just above), at density 10^5^ cells.mL^−1^, with two replicates each. Calibration samples were frozen, the chloroplast and ITS2 loci were amplified, sequenced, and all reads converted to short haplotypes as described above. No *viridis* alleles were detected at the chloroplast locus where primers were specifically designed to amplify our strains, and we observed a correct match between observed and expected frequency (adjusted *r*^2^ = 0.89). Given that such correlation achieved in the presence of contaminants is extremely likely to hold in the absence of *viridis* contamination in the analysis of the experimental populations, we use the allele C frequency measured at the chloroplast locus as a direct proxy for strain C frequency. At the ITS2 locus, after removal of all *viridis* sequences, we found a perfect linear relationship between the frequencies of allele C *p*_ITS2_ and the strain C frequency *p* (*p_ITS_*_2_ = 0.19 + 0.79*p*, adjusted *r*^2^ > 0.99).

#### State-space logistic regression

We estimated fluctuating selection by tracking the dynamics of the frequency *p* of strain C through time. Population genetic change under selection (especially fluctuating selection) is more conveniently analyzed on the logit scale, where responses to selection are additive over time (Chevin, 2011, 2019; Gillespie, 1991 p147; Kimura, 1954). The logit is also the canonical link function for a generalized linear model with binomial error (logistic regression), which is well-suited for population genetic measurements of selection (Gallet et al., 2012).

Here we estimated strain frequencies by combining two sources of genetic information, from the ITS2 and the chloroplast locus. Our rationale was that if the two loci recombine very little − as confirmed by the strong linear relationship between allelic frequencies at both loci (*r*^2^ = 0.96), which remained unchanged over time (P = 0.27 for time effect on the regression slope of allele C frequency measured at the ITS2 against at the chloroplast locus) −, then using both as indicators of strain identity makes more efficient use of the data than performing simple, univariate logistic regression on each marker. We thus considered that the measured frequencies at the ITS2 and chloroplast loci were two observations (with error) of a true, unobserved strain frequency *p*. Formally, this corresponds to a state-space model (Kedem & Fokianos, 2002), where the true dynamics of strain frequency is treated as an unobserved underlying process, while the observations (frequencies at the ITS2 and chloroplast locus) have errors that are mutually independent after conditioning by the underlying process. A state-space model is thus fully specified by the distributions of the errors and the underlying process. We wrote an explicit likelihood function in C++, and optimized it using the TMB package in R (v.3.5.2). R and C++ codes are available from a digital repository.

#### Observation model

At the chloroplast locus, the number *n_i,t_* of C sequences in population *i* at time *t* was assumed to follow a binomial distribution with parameters the strain frequency *p_i,t_* and the total number of chloroplast sequences 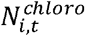. At the ITS2 locus, the number *m_t_* of sequences C followed a similar binomial distribution, with a linear correction (with coefficients *α* and *β*) to account for the presence of a logit frequency *α* of allele C in strain A:

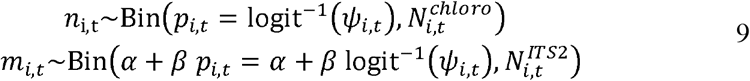

### Process model

In each population *i*, the dynamics of allele C frequency *p* are such that:

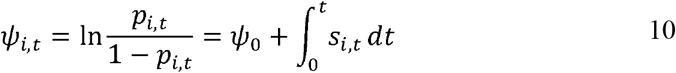

Where *ψ_i,t_* is the logit frequency of allele C, and *s_i,t_* the selection coefficient in population *i* at time *t.* In other words, selection coefficients are integrated/summed over time in their contribution to logit allelic frequency *ψ*.

Integration of the stochastic differential equation (10) leads to the distribution of allele C logit frequency at time *t.* In particular, when *s_i,t_* follows a Gaussian process, then logit frequency also has a Gaussian distribution, with mean and variance that increase linearly over time, with slopes 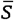 (the mean selection coefficient), and 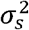 (the variance of selection coefficients), respectively (Gillespie, 1973, 1977). More generally, summing/integrating selection coefficients over time in eq. (10) should eventually lead to a normal distribution of *ψ_i,t_* because of the central limit theorem, so we used a Gaussian process to model the dynamics of logit frequency,

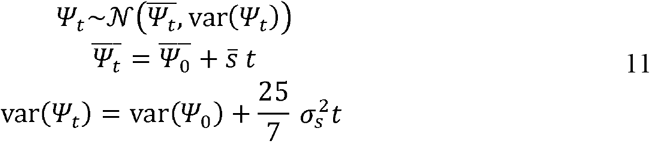

The factor 25/7 in eq. (11) serves as a correction for the fact that selection does not fluctuate continuously over time, but instead remains constant during the time interval between our bi-weekly transfers, which affects estimation of the variance but not the mean of selection (Appendix). We investigated the influence of our experimental treatments on these parameters of fluctuating selection, by including the environmental mean *µ*^2^*_E_* (corresponding to the deviation from the mean salinity 2.4M), squared mean *µ^2^_E_*, variance 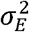, and predictability 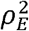, as covariates for 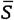and 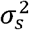 in the regression in eq (11), leading to

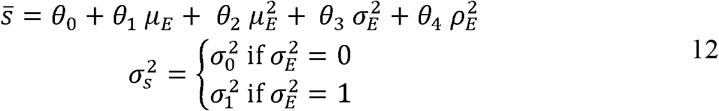

where *θ*_0_ is the selection coefficient in constant 2.4M salinity, and *θ*_1_ to *θ*_4_ are the coefficients associated with the mean, squared mean, variance and predictability of the environment, respectively.

We then searched for finer, short-scale mechanisms underlying these aggregate macroscopic effects of fluctuating selection, by focusing on frequency change over subsequent transfers with known salinities. We tested for effects on selection of current and previous salinity (following the reaction norms in eqs 2 and 5), as well initial frequency, and initial population density. For each transfer, frequencies both before and after selection were considered as random variables, and estimated from the observation model.

Combining the quadratic selection reaction norm in eq. (2), estimated over the long run in constant environments, with the normal distribution of environments in our fluctuating treatments, leads to analytically tractable distributions of predicted selection coefficients in a fluctuating environment. When the selection reaction norm is linear (c = 0), the distribution of selection coefficients becomes normal (**Figure 5. b** and f), as expected when a mutation does not cause differences in plasticity or tolerance breadth between genotypes (Chevin, 2019). In contrast, a concave selection reaction norm with c ≠ 0 leads to a displaced non-central chi-square distribution (with one degree of freedom) for *s*. This distribution may be highly asymmetric if its non-centrality parameter is small, that is, if the linear term is small relative to the quadratic term in eq. (2). This occurs when the average environment is close to the optimum environment for selection (**Figure 5 d, e** and f).

## Acknowledgements

We thank P. Nosil and C. Leung for useful comments on this manuscript. This work was supported by the European Research Council Grant 678140 (FluctEvol).

## Supporting information

### Online Appendix

Here, we investigate how our experimental set up consisting of successive transfers of 3 and 4 days, with constant salinity in between transfers, impacts the stochastic dynamics of allelic frequencies.

Between transfers *T –* 1 and *T*, each population *i* of a fluctuating salinity treatment grows in a specific salinity *E_i_*_,*T*_, which remains constant until the next transfer. Assuming frequency-and density-independent selection, the selection coefficient *s_i_*_,*T*_ is also constant for each line in between two transfers (and more generally, even when the selection coefficient does vary, we can only estimate an average selection coefficient *s_i_*_,*T*_ over the interval between two transfers). The logit-transformed allelic frequency in line *i* just before the next transfer is thus (from eq. 10 in the main text):

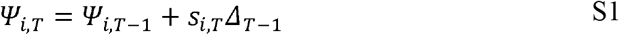

where Δ*_T–1_* is the time interval between transfers *T –* 1 and *T*. Assuming a normal distribution of distribution coefficients (with mean 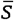 and variance 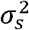), the distribution of allelic frequencies in the *n* lines under the stochastic treatment is also normal, with mean

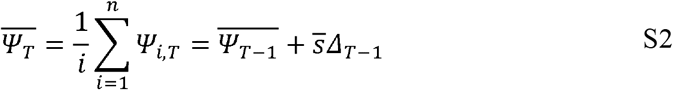

and variance:

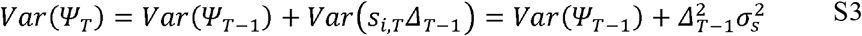

Note that the variance does not increase linearly, but quadratically with the time interval between transfers Δ*_T–1_*.

After the next transfer, the mean and variance of logit allele frequency are:

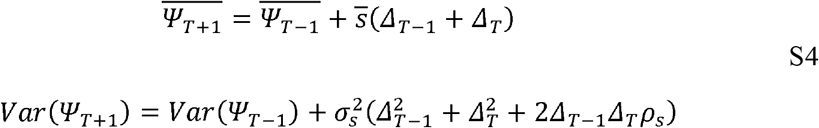

where *ρ_S_* is the correlation of selection coefficients between subsequent salinities in the time series. In our experimental set up, we had bi-weekly transfers, such that Δ*_T–1_* = 3 (or 4) and Δ*_T_* = 4 (or 3), leading to:

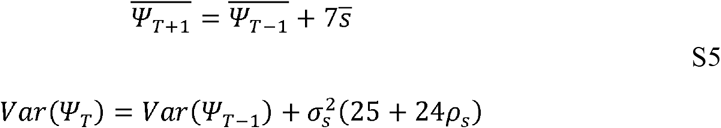

In the logistic regression described in the main text, we infer the mean and variance of selection as the slopes of temporal changes in the mean and variance of logit allelic frequencies, with unit one day. After two transfers, (from *T* – 1 *T* + 1), we thus estimated the mean and variance of selection as:

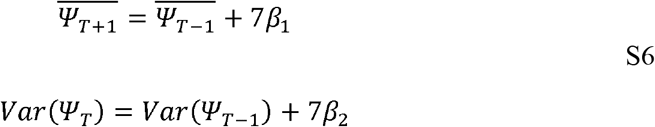

For the mean selection coefficient, comparing eqs. S5 and S6, we directly have 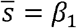. For the variance of selection coefficients, neglecting autocorrelation *ρ_S_* for simplicity, we get 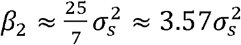. This means that because of our experimental set up where salinity remains constant in between transfers, the variance of logit frequency increases faster with time than in a true discrete-time or continuous-time stochastic set up with the same duration. Therefore, to directly estimate 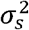 in our logistic regression framework, we multiply the coefficient in the regression for the variance by 25/7 in eq. (1).

## Supplementary Tables

**Supplementary Table 1:** Effect of past (*E*_t-1_) and current (*E*_t_) environment on selection (Estimates, Standard errors and P-values from Wald test).

|  | *Estimate* | *Std. Error* | *Pr(>\|W\|)* |
| --- | --- | --- | --- |
| 1 | 0.0437 | 0.02451 | 7.48E-02 |
| $E_t$ | 0.0775 | 0.0198 | 9.33E-05 |
| $E_t^2$ | -0.004528 | 0.0174 | 7.95E-01 |
| $E_{t-1}$ | -0.0400 | 0.0196 | 4.12E-02 |
| $E_{t-1}^2$ | 0.0267 | 0.0181 | 1.39E-01 |
| $E_{t-1} * E_t$ | 0.0312 | 0.0181 | 8.50E-02 |
| $\sigma_s^2$ | 0.00521 | 0.000594 | 1.80E-18 |

## Supplementary Figures

**Supplementary Figure 1:**
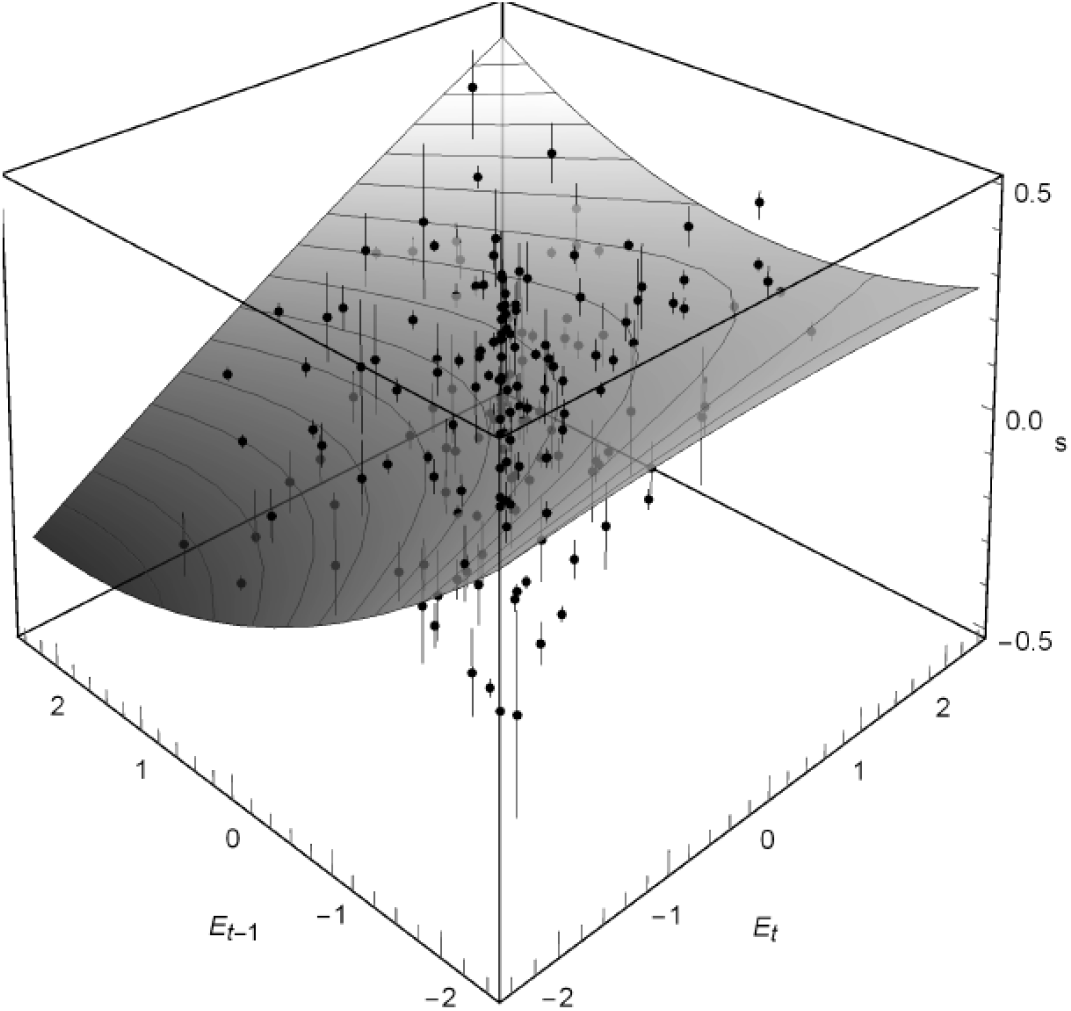
Selection reaction norm (*s*) fitted during on transfer depends on past (*E_t-1_*) and current (*E_t_*) environments. Selection coefficients and their standard errors (dots and error bars) were computed from the realized logit allele frequencies estimated at two successive transfers.

**Supplementary Figure 2:**
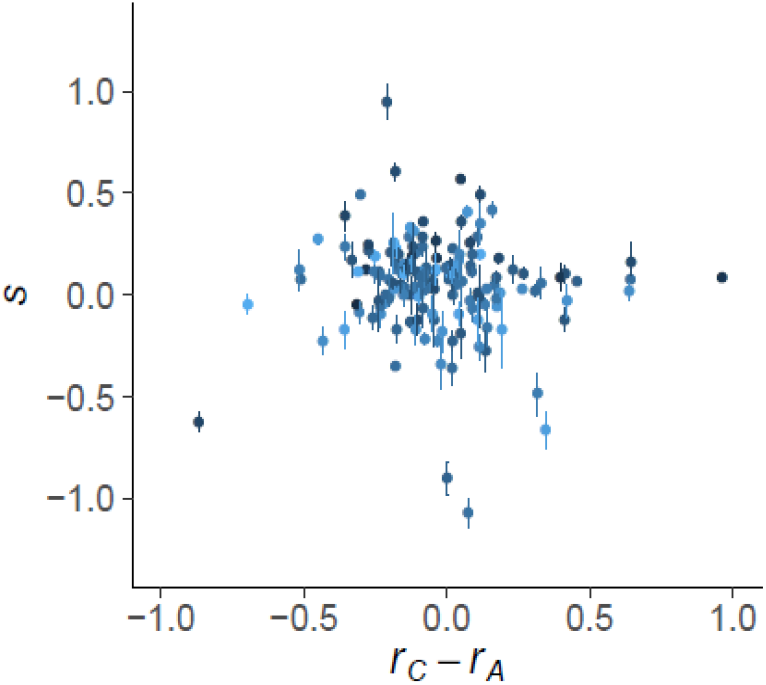
Difference between growth rate in isolation of strain C and A (x) does not correlate with selection coefficients estimated between two successive transfers (y). Growth rates estimates were extracted from Rescan et al. (2020), and selection coefficients were computed from the realized logit allele frequencies estimated between two successive transfers. Colors correspond to the salinity before transfer, from blue (0 M) to black (4.8 M)

## Notes

### Competing Interest Statement

The authors have declared no competing interest.

